# Co-evolution integrated deep learning framework for variants generation and fitness prediction

**DOI:** 10.1101/2023.01.28.526023

**Authors:** Xiaoqin Tan

## Abstract

Pandemic caused by viral protein is characterized by waves of transmission triggered by new variants replacing old ones, resulting in immune escape and threatening public health. Therefore, there is an obvious need to accurately identify the vital mutation sites and understand the complex patterns of mutation effect of viral protein. However, existing work do not explicitly modelling vital positions functioning for virus fitness, leading to large search space with money- and time-consuming search cost. Here, we propose EVPMM (evolutionary integrated viral protein mutation machine), a co-evolution profiles integrated deep learning framework for dominant variants forecasting, vital mutation sites prediction and fitness landscape depicting. It consists of a position detector to directly detect the functional positions as well as a mutant predictor to depict fitness landscape. Moreover, pairwise dependencies between residues obtained by a Markov Random Field are also incorporated to promote reasonable variant generation. We show that EVPMM significantly outperforms existing machine learning algorithms on mutation position detection, residue prediction and fitness prediction accuracies. Remarkably, there is a highly agreement between positions identified by our method with current variants of concern and provides some new mutation pattern hypothesis. The method can prioritize mutations as they emerge for public health concern.

## 1 Introduction

Viral protein is highly mutable, such as influenza[1], HIV[2] and SARS-CoV-2[3, 4], resulting pandemic by repeated waves of cases by the emergence of new variants with higher fitness, which fitness refers to the ability of viral proteins to replicate, propagate and infect[5]. Mutations observed in prevalent variants of interest (VOIs) or variants of concern (VOCs) are associated with increased transmissibility and reduce efficacy of antibody and immune response[6]. Therefore, rapid and accurate identification of vital mutation sites and forecast dominant variants is critical for guiding outbreak response. Moreover, the knowledge of the fitness landscape of viral proteins plays a key role in the rational design of vaccines.

There are several machine learning methods in recent years, to modelling diversity of proteins and the corresponding variants mutation effect can be scored. But most of these methods are not specific to viral protein, leading to ambiguous prediction due to scarce viral protein dataset. In addition, existing methods are unable to capture key mutation sites that are critical for fitness. Therefore, we argue that current machine learning models for reasonable virus variant generation have the following three limitations: (1) do not pay much attention to the modeling of mutational positions. Mutational positions are conserved sites in viral protein, which influence vieal protein function and folding. Besides, mutation positions are extremely sparse in viral protein sequences. The average protein length is 500 amino acids and even 1273 for SARS-CoV-2 spike protein while less than 10% positions functional for virus fitness. Thus, explicitly modeling critical positions can obviously reduce the enormous search space and also save time.(2) Epistasis effect is universal, and combined mutations are far greater than single point mutations in the real world. None of these machine learning methods such as CSCS[7], EVmutation[8], DeepSequence [9] are tailored for combinatorial mutations. (3) Few methods are discussed the relation about model and the higher-order dependencies of residues.

Due to the importance of vital mutation sites identification, we propose EVPMM (evolutionary integrated viral protein mutation machine) to explicitly detect vital mutation positions, forecast variants with higher fitness and depict the fitness landscape of variants (Figure 1). Specifically, EVPMM consists of a position detector and a mutant predictor. The position detector has the ability to directly find the vital positions functioning for virus fitness, thus obviously reducing the search space. Based on the detected positions, mutant predictor can recognize the favorable amino acid under the given fitness feature at a specific position by leveraging (a) the contextual information of viral protein sequence from the neural network and (b) the global pairwise dependencies between residues from multiple sequence alignment (MSA).

**Fig. 1.**
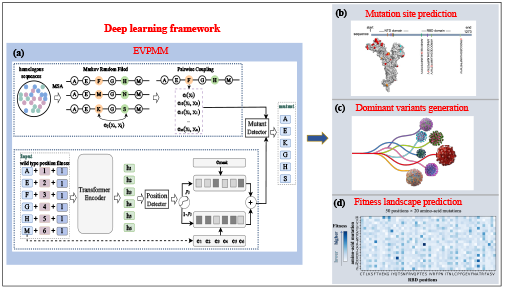
Overview of EVPMM. (a) EVPMM consists of a position detector and a mutant predictor. We assume the wildtype sequence is (AEFGHM) and the mutant sequence is (AEKGHS), where the mutation occurs at position 3 and 6. emask denotes the embedding of [mask] symbol and e* denotes the token embedding at the corresponding position. A deeper color in emask and e* represents a higher probability.(b) EVPMM can accurately detect the mutation position. (c) The model can generate variants with higher fitness. (d) The model can depict the landscape of variants.

Thus, the whole framework is capable of detecting vital positions with a low search cost and generate variants with higher fitness to improve health care in advance. Also, the model can provide fitness landscape of predicted variants, contribute to the rational design of vaccines. Experimental results on three viral protein datasets show that EVPMM significantly outperforms several strong baselines on mutation position detection, residue prediction and fitness prediction accuracies. Besides, we also find that there is an highly agreement between the mutational positions detected by EVPMM and those in VOCs and VOIs such as Omicron BA.1, demonstrating EVPMM can accurately detect dominant mutation positions with a low search space. Remarkably, our model also discovers a potential functional position 487 of SARS-CoV-2 that has not been reported by any other work, at which the generated variant may bind better with ACE2. These results demonstrates that EVPMM can be used as an useful tool to detect vital mutation positions and forecast variants with higher fitness.

## 2 Results

### 2.1 Architecture of EVPMM

We propose a novel neural network model called EVPMM to generate virus variants, as illustrated in Figure 1. Given a wild type X = {x1, x2, …, xn} and its fitness label *Z*, EVPMM targets at generating a new variant Y = {y1, y2, …, yn} that expresses the corresponding fitness feature, *e*.*g*. P(Y ‖ X, Z).n is the number of amino acids in X and Y. EVPMM consists of a position detector and a mutant predictor. Position detector aims to detect the possible positions that are likely to mutate into a different amino acid, while the mutant predictor targets at predicting the mutated amino acids given the specific mutational positions. The overall training objective can be formulated as *L* = *L*pos + *λL*mutew, where *L*pos denotes the position detection loss and *L*mut for amino acid prediction loss. The importance of the two losses are controlled by a hyperparameter *λ*. Through this process, EVPMM can find the potential virus variants that have higher fitness advantage in nature.

Position detector is designed to directly find the functional positions which possibly play an important role in protein structure stability and virus infection. At a specific positon *i*, the model input is composed of token embedding Emb(*xi*), position embedding Emb(*posi*) as well as fitness label embedding Emb(*Z*). Thus, the input contains the residual type, sequence position information, and is conditioned on fitness. Then the informative embedding was as input to the encoder layer of EVPMM. Encoder layer contains *L* Transformer [10] encoder layers, and each layer consists of a self-attention block followed by a position-wise feed-forward block. Using the position-wise feed-forward block, the model provides the probability distribution of each specific position. The 2-D one-hot vector denoting if the amino acid at position *i* is changed while 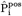 is the predicted one, and 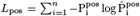 as the loss to detect the mutable position.

Mutant predictor is designed to obtain the priority amino acid at a predicted mutated position. The mutated amino aicd probability distribution is composed of two parts, respectively from viral protein sequence and multiple sequence alignments (MSA). At this section, the model integrated evolutionary profile to model the residue dependencies in protein sequences as 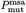, in which a Markov random field (MRF) model was established on MSA profile to capture epistatic effect. Through the position detector, we get the mutation positions and the probability distribution of each mutated position, then through the weight assignment as as 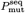, emask denotes the embedding of [mask] symbol and e* denotes the token embedding at the corresponding position. Then the mutation predictor probability at a specific position is 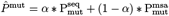, then 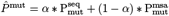 as the loss of mutant predictor.

### 2.2 EVPMM can accurately detect the mutation positions

Viral protein can evolve variants that abrogate the binding of antibodies or T cell receptors,while also preserving or even increasing fitness. To accurate the vital mutations of the next waves of variants is urgently in need for world health care protection. Thus, the accuracies of the mutation positions detection is particularly important.

To indicate the accurate of EVPMM on mutation positions detection, we compare our proposed EVPMM against six representative baselines, including statistical methods and neural network models. We compare our proposed EVPMM against the following representative baselines, including statistical methods and neural network models.

Statistical methods explicitly model the co-variations between all pairs of residues in the protein by fitting a statistical model to the multiple sequence alignment (MSA) of all homologous sequences of the protein of interest: (1) MAFFT MSA [11] uses an algorithm based on progressive alignment to create MSA of amino acid sequence. (2) EVMutation [8] is an unsupervised statistical model which fits a pairwise undirected graphical model to the multiple sequence alignment.

The following neural network models are also taken as strong baselines: (3) CSCS proposes a LSTM-based language model to learn the mutational semantics of viral proteins. (4) Trans-MLM: We also realize a Transformer-Encoder [10] based language model as CSCS, which is optimized using masked language model (MLM) task. We call it Trans-MLM. (5) Trans-Mut: We also realize a Transformer-Encoder [10] called Trans-Mut which takes wild type as input and virus mutant as output. (6) DeepSequence [9] is a generative model that predicts the effects of mutations based on variational inference [12, 13] to capture higher-order dependencies of residues in the protein.

EVPMM is better at predicting a favorable amino acid under the given fitness feature at a specific position. As shown in Figure 2, EVPMM achieves the highest normalized AUC score on all three virus datasets. Our interpretation is that the amino acid prediction is realized by incorporating both contextualized viral protein information as well as global pairwise correlation from MSA, which benefit better variant prediction. In most cases, neural network models performs better than statistical methods, as deeper features captured by neural networks are helpful for virus mutant prediction. CSCS achieves the worst performance among the five neural network models, demonstrating that Transformer-encoder based model is better at viral variant prediction than LSTM-based model.

**Fig. 2.**
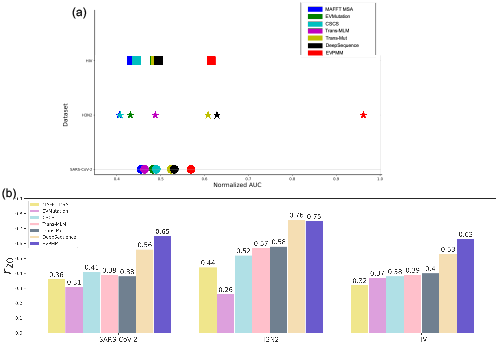
Comparison to the six strong baselines on SARS-CoV-2, H3N2, HIV dataset. (a) AUC: We evaluate the accuracy of amino acid prediction using AUC which is obtained by plotting acquired fitness mutations versus total acquired mutations based on 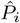, normalized by the maximum area to produce a score between 0 and 1.(b) *r*20: *r*20 is proposed by [14] to evaluate the generation capability of protein generative models by capturing the higher-order dependencies of residues.

Pairwise correlation is beneficial for virus mutant prediction. The AUC and *r*20 score both decrease significantly after removing the pairwise dependencies though removing position detector hurts performance more. It demonstrates that involving dependencies of residues in viral protein sequence is beneficial for capturing higher-order correlations, which are helpful to generate reasonable virus variants.

To better understand the mutational effect, we count the mutational position overlaps between the generated virus variant by our EVPMM and the existing SARS-CoV-2 variantshttps://viralzone.expasy.org/9556. The results are shown in Table 1. It shows that there exists much overlap between the mutational positions detected by our EVPMM and those reported in other variants, demonstrating that the virus sequence generated by our model may have high virus fitness due to the similarity to the existing variants.

**Table 1.** Mutational position overlap between the generated samples of the proposed EVPMM and the existing SARS-CoV-2 variants.

| Position | AA Wuhan-Hu-1 | Pango Lineage |
| --- | --- | --- |
| 417/501 | K/N | B.1.351, P.1, B.1.1.28.1, Omicron BA.1, Omicron BA.2, RBD-ACE2 |
| 484 | E | B.1.351, P.1, B.1.1.28.1, Omicron BA.1, Omicron BA.2 |
| 339/371/373 | G/S/S | Omicron BA.1, Omicron BA.2 |
| 375/478/498 | S/T/Q | Omicron BA.1, Omicron BA.2 |
| 367 | V | A.23.1 |
| 452 | L | B.1.617.2, B.1.427, B.1.429 |
| 446 | G | RBD-ACE2, Omicron BA.1 |
| 496/547 | G/T | Omicron BA.1 |
| 376 | T | Omicron BA.2 |
| 453 | Y | RBD-ACE2, B.1.1.298 |
| 449/500 | Y/Y | RBD-ACE2 |

### 2.3 EVPMM can accurately predict the functional fitness of viral protein

Fitness refers to the relationship of sequence and function. When introducing novel mutation, the function of viral protein is changed, resulting the fitness of viral protein increase or decrease. Thus, accurately predict the fitness is vital to variants detection and world health care protection in advance. Also, the knowledge of the fitness landscape of viral proteins plays a key role in the rational design of vaccines.

Here, we compare EVPMM to state of the art of model DeepSequence on six HA datasets. As shown in Figure 3, EVPMM outperformes DeepSequence on BK79, Bris07L194, Bei89, NDako16, Mos99 and all HA datasets. The results indicate EVPMM has a strong ability on fitness prediction on different viral variants. All these HA datasets are combinational mutation dataset, the results indicate that EVPMM has a strong ability to capture combinatorial mutation pattern than existing SOTA model.

**Fig. 3.**
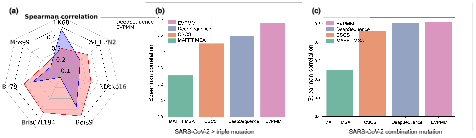
Comparison to the strong baselines on SARS-CoV-2, HA dataset on fitness prediction. (a) Prediction the fitness of six HA datasets. The result predicted by EVPMM is positively correlated with fitness, outperforms SOTA model DeepSequence.(b) Comparison to strong baselines on SARS-CoV-2 datasets.

As shown in Figure 3, EVPMM can accurately predict fitnss on combinational datasets. SARS-CoV-2 datasets contain lots of single and combinational mutation. On single, double and all combinational mutation datasets, EVPMM has a comparable performance to DeepSequence. But on the multiple sites mutation dataset that mutation positions count greater than three, EVPMM outperformed DeepSequence and other strong baselines. These results demonstrate EVPMM has a strong ability to fitness prediction, especially to capture combinational mutation patterns.

### 2.4 EVPMM provides rational fitness landscape

Fitness landscape is informational to rational vaccine design and protein functional research. EVPMM provide rational fitness landscape on different mutated sequences. As shown in Figure 3, the landscape predicted by EVPMM has a highly aggrement to experimental fitness landscape on SARS-CoV-2. These results indicate EVPMM provides rational priority amino acids distribution at a specific position. We first visualize the mutational probability distribution of each position in the wild type sequence of SARS-CoV-2, and discover that the mutations focus on positions 330 ∼ 380, as illsutrated in Figure 4.

**Fig. 4.**
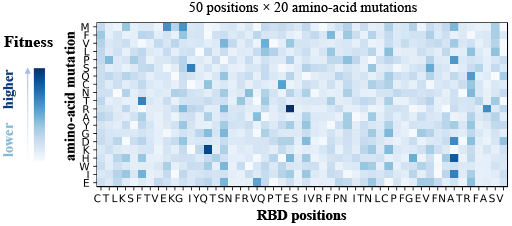
The mutational effects of positions 330 380 in the wild type sequence of SARS-CoV-2, where the mutations occur most.

### 2.5 Case study

To gain an insight on how well the virus variants are generated through our proposed EVPMM, we visualize some generated SARS-CoV-2 cases. First, EVPMM generates some variants through sampling method. Then we visualize the produced samples using PyMOL tool [15], and the results are illustrated in Figure 5. Figure 5(a) illustrates the generated SARS-CoV-2 variants are highly enriched in the binding site between ACE2 and RBD block, demonstrating our EVPMM can precisely predict mutable positions of the virus. Figure 5(b) illustrates that the generated variant may have an optimal binding with ACE2 at a potential mutational position 487, which has not been reported by any other work, validating that EVPMM has the ability to produce diverse and novel SARS-CoV-2 variants.

**Fig. 5.**
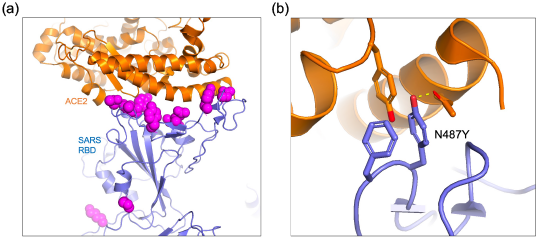
3D visualization of (a) generated sample binding with ACE2, (b) new position detected by our model.

### 2.6 Ablation study

To better analyze the influence of different components in our EVPMM, we conduct ablation tests and the results are shown in Table 3.

Position detection network is critical for combinatorial position prediction.After removing position detector, the detection accuracy drops significantly. It validates that the position detector plays an important role in vital position recognition, which functions for virus fitness, such as structure stability, virus infection as well as transmissibility.

Pairwise correlation is beneficial for virus mutant prediction. The AUC and *r*20 score both decrease significantly after removing the pairwise dependencies though removing position detector hurts performance more. It demonstrates that involving dependencies of residues in viral protein sequence is beneficial for capturing higher-order correlations, which are helpful to generate reasonable virus variants. We also test if the well-pretrained protein Bert can further improve the performance of our model. We take the 6-layer ESM model [16] as initialization https://github.com/facebookresearch/esm, and the results are shown in Table 2. From the table, we can see that finetuning from ESM can benefit model performance on H3N2 datset, but it degrades performance on SARS-CoV-2 and HIV-1 datasets. On one hand, ESM adopts masked language modeling task which is different from our training paradigm. On the other hand, our model distinguish training data with different fitness preference while ESM does not take this into account. We guess these differences in training lead to performance unimprovement. However, pretraining may help model summarize information from large-scale protein data, and thus it can benefit model from some aspects.

**Table 2.** Results of EVPMM finetuned from existing protein Bert (ESM).

| Models | SARS-CoV-2 |  |  | H3N2 |  |  | HIV-1 |  |  |
| --- | --- | --- | --- | --- | --- | --- | --- | --- | --- |
| | AUC | Detection | $r_{20}$ | AUC | Detection | $r_{20}$ | AUC | Detection | $r_{20}$ |
| EVPMM | **0.569** | **0.999** | **0.65** | 0.962 | **0.997** | **0.75** | **0.615** | 0.995 | **0.63** |
| – ESM finetuning | 0.555 | 0.998 | 0.58 | **0.963** | **0.997** | 0.74 | 0.549 | **0.998** | 0.61 |

## 3 Related Work

### 3.1 Viral Variant Generation

When in-depth knowledge of protein structure and phenotype, some physical-based methods [17, 18] such as FoldX [19], PoPMuSiC [20] and IMutant [21] can be used to putative protein variants. Applicable more generally, statistical models take advantage of evolutionary profiles based on MSA to capture second-order co-evolution for functional viral protein design. Rodriguez-Rivas *etc*. propose an energy-based probabilistic graphical model to capture position dependencies and infer unseen risky variants of SARS-CoV-2 [6]. These probabilistic models have also been applied in HIV [22, 23], poliovirus [24] and influenza [25]. However, these statistical models are limited to pairwise epistatic interactions and ignore high-order covariation patterns, while some evidence suggests that higher-order epistasis affecting protein evolution.

Some machine learning based models are developed to capture higher-order interactions for viral protein variants generation [9, 26–29]. VPRE is a VAE-based model that utilizes real-time and simulated SARS-CoV-2 genomic data to learn a continuous latent space to model the trajectory the SARS-CoV-2 spike protein evolution [30]. MutaGAN is a framework using GANs with RNNs to generate complete viral protein sequences augmented with possible mutations of future virus populations [28]. Furthermore, multiple language-based models were proposed [31], to capture the functional effect of sequence variation by training on naturally viral-specific sequences. Overall, these models do not consider any position information and they do not truly capture higher-order epistasis, but rather benefit from biologically motivated priors and engineering efforts [14, 29].

#### 3.1.1 Functional Protein Design

Functional protein design has attracted a lot of research interest recently. Currently computational methods can be divided into three categories. The first is physical-based methods [32–34], which use Monte Carlo simulation to modify the sequence or its structure iteratively to achieve a local energy minimization. The second is sequence-based methods [35–37]. For example, [38] formulate protein design as an autoregressive process to generate a diverse sequences library for nanobody. The third is 3D-structure based functional protein design [39–41]. Fold2Seq was proposed as a novel transformer-based generative framework for designing protein sequences conditioned on a specific target fold [40]. A generative model was proposed to specifically design the functional motif of antibodies, by co-designing the sequence and 3D structure of CDR [41].

To better capture the mapping of protein sequence and function for further functional protein design, a lot of machine learning based methods have been developed to predict mutation effect [8, 9, 42–44]. Models were developed for protein fitness landscape prediction, including feature engineering models such as Envision [45], language models such as TAPE [46], ESM-1v [47], MSA Transformer [48], ensemble models such as CADD [49], M-CAP [50], Revel [51], statistical methods [8, 52, 53] and representation learning methods [54, 55]. Our proposed method explicitly captures combinational mutation patterns on the one hand and learns the fitness landscape for mutational position on the other, providing insights into the relationship between viral protein evolution and function.

## 4 Methods

In this section, we describe our proposed EVPMM in detail, which aims to generate diverse and reasonable virus variants in order to detect potential variants in the future and take action in advance. In the following subsections, we will first introduce the overall architecture of our model. Then we describe how to directly find the functional positions, based on which we will introduce how to predict the mutated amino acids with higher fitness. Finally, a pivot-based data augmentation method is proposed to further improve the diversity of the virus data.

### 4.1 Overall Architecture

We propose a novel neural network model called EVPMM to generate virus variants, as illustrated in Figure 6.

**Fig. 6.**
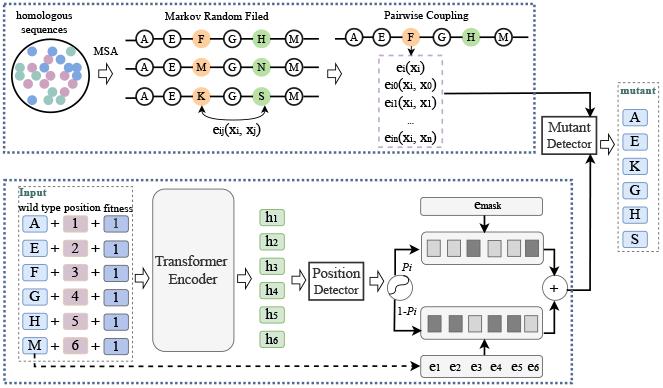
The architecture of the viral protein mutation machine (EVPMM). We assume the wild type sequence is (AEFGHM) and the mutant sequence is (AEKGHS), where the mutation occurs at position 3 and 6. emask denotes the embedding of [mask] symbol and e* denotes the token embedding at the corresponding position. A deeper color in emask and e* represents a higher probability.

Given a wild type X = {x1, x2, …, xn} and its fitness label *Z*, EVPMM targets at generating a new variant Y = {y1, y2, …, yn} that expresses the corresponding fitness feature, *e*.*g*. P(Y ‖ X, Z).*n* is the number of amino acids in X and Y. EVPMM consists of a position detector and a mutant predictor. Position detector aims to detect the possible positions that are likely to mutate into a different amino acid, while the mutant predictor targets at predicting the mutated amino acids given the specific mutational positions. The overall training objective can be formulated as follows:

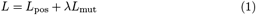

where *L*pos denotes the position detection loss and *L*mut for amino acid prediction loss. The importance of the two losses are controlled by a hyperparameter *λ*. Through this process, EVPMM can find the potential virus variants that have higher fitness advantage in nature.

### 4.2 Position Detector

Position detector is designed to directly find the functional positions which possibly play an important role in protein structure stability and virus infection. Specifically, EVPMM is composed of *L* Transformer [10] encoder layers, and each layer consists of a self-attention block followed by a position-wise feed-forward block, which can be defined as follows:

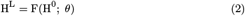

where 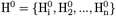 and 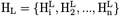 respectively denotes the model input and *L*-th layer output. *θ* are the trainable model parameters. At a specific positon *i*, the model input is composed of token embedding Emb(*xi*), position embedding Emb(*posi*) as well as fitness label embedding Emb(*Z*):

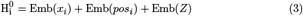

Finally, the position detector predicts whether a position is mutable under the given fitness feature as follows:

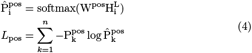

where W^pos^ is the trainable model parameter. 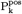 is a 2-D one-hot vector denotes if the amino acid at position *k* is changed while 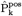 is the predicted one.

Using the position detection network, EVPMM has the capability to directly capture the combinatorial positions that may have higher structure stability, virus infection or transmissibility, which can largely reduce the search space.

### 4.3 Mutant Predictor

Through the position detector, a subset ℙ𝕊 is gathered representing the possible functional positions, based on which the mutant predictor aims to predict the possibly mutated amino acids. The mutated amino aicd probability distribution is composed of two parts, respectively from viral protein sequence and multiple sequence alignments, which are controlled by a hyperparameter *α*. Thus, the mutation predcition loss can be defined as:

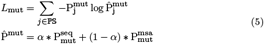

where 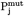 is a 20-D one-hot vector denoting the mutated amino acid at position *j*.

To calculate the distribution from viral protein sequence, gumbelsoftmax [56] is leveraged:

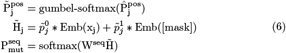

where 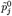 and 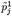 respectively denotes the 0- and 1-dimensional value of 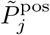. Emb([mask]) denotes the embedding of [mask] symbol. Then, to better capture high-order amino acid covariations, we apply CCMpred [57, 58], which is based on a Markov random field (MRF) specification to model the residue dependencies in protein sequences. The energy function E(X) of sequence X is defined as the sum of all pairwise coupling constraints *eij* and single-site constraints *ei*, where i and j are position indices along the protein sequence:

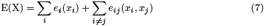

When the MRF model is fit to data with proper regularizations, the residue dependencies in protein sequences are explained by the direct coupling terms *eij*. Finally, the probability distribution from pairwise coupling can be computed as follows:

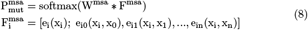

where W* denote trainable parameters.

Through this process, the amino acid prediction is calculated considering both contextual information from protein sequence as well as global coupling from MSA, which benefits better virus variant generation.

### 4.4 Pivot-Based Data Augmentation

To improve the diversity of virus data, we propose a pivot-based data augmentation method, which is inspired by pivot-based machine translation [59] in natural language processing. Specifically, one wild type could have many mutational variants in nature, each of which has a corresponding fitness value.

To increase the diversity of <wild type >, variant> pairs, we make the following rule: if the fitness value of variant A is lower than variant B, then we think B is more adaptable to the environment than A and give pair <A, B>with fitness label 1 otherwise 0. Thus, pair <A, B >with label 1 represents that A mutates into B with a higher fitness advantage such as structure stability, binding to receptors or transmissibility, and vice versa. Through this way, abundant data pairs can be produced, which are helpful for diverse virus variant generation.

### 4.5 Datasets and training details

#### Datasets

We evaluate our model on three viral protein datasets, including H3N2 causal fitness dataset, HIV-1 Envelope (Env) protein dataset as well as SARS-CoV-2 viral protein dataset [7]. We use all the fitness part of these datasets, which are composed of combinatorial mutations. All the fitness values are averaged, and the pair of which the fitness value is larger than average is given label 1 otherwise 0. we randomly split each dataset into training/validation/test sets with the ratio of 8:1:1 (9:0.5:0.5 for SARS-CoV-2). For each pair in the training split, we randomly sample a number *N* from a uniform distribution with minimum value 5 and maximum value 50, and then construct *N* pseudo pairs using our proposed pivot-based data augmentation method. The detailed statistics are shown in Table 4.

**Table 3.** Ablation study results on three viral protein datasets. Detection denotes the position detection accuracy.

| Models | SARS-CoV-2 |  |  | H3N2 |  |  | HIV-1 |  |  |
| --- | --- | --- | --- | --- | --- | --- | --- | --- | --- |
| | AUC | Detection | $r_{20}$ | AUC | Detection | $r_{20}$ | AUC | Detection | $r_{20}$ |
| EVPMM | **0.569** | **0.999** | **0.65** | **0.962** | **0.997** | **0.75** | **0.615** | **0.995** | **0.63** |
| – w/o Pos | 0.541 | 0.421 | 0.61 | 0.684 | 0.848 | 0.67 | 0.491 | 0.134 | 0.51 |
| – w/o MSA | 0.564 | 0.979 | 0.60 | 0.821 | 0.991 | 0.71 | 0.602 | 0.923 | 0.56 |
| – retrieve | 0.531 | 0.783 | 0.57 | 0.583 | 0.875 | 0.63 | 0.472 | 0.637 | 0.49 |

**Table 4.** Statistics of viral protein datasets.

| Datasets | Training | Validation | Test | Pivot-Based Augmentation | Length |
| --- | --- | --- | --- | --- | --- |
| H3N2 | 2761 | 345 | 345 | 119166 | 560 |
| HIV-1 | 20248 | 2531 | 2531 | 832648 | 852/860 |
| SARS-CoV-2 | 95526 | 5000 | 5000 | 633156 | 1273 |

#### Experimental Settings

EVPMM is constructed based on Transformer [10] encoder with 6 layers. The model hidden size *dmodel* and feedforward hidden size *dff* are set to 512 and 2048 respectively. The number of attention head is set to 8. The vocabulary size is set to 20 with common amino acids. The mini-batch size and learning rate are set to 64k tokens and 7*e* − 5 respectively. The model is trained with 8 NVIDIA Tesla V100 GPU cards and the maximum training step is 300, 000. We apply Adam algorithm [60] as the optimizer with a linear warm-up over the first 10, 000 steps and linear decay for later steps, of which the initialized and maximum learning rate are set to 1e-8 and 1e-11 respectively. We tune the hyper-parameter *λ* and *α* both from 0.1 to 1.0 with step 0.1, and find that *λ* = 0.5 and *α* = 0.7 achieves the minimum loss on the validation set.

## 5 Conclusion

This paper proposes EVPMM, a viral protein variant generation model. It is composed of a position detector and a mutant predictor. The position detector targets at directly finding the functional positions while the mutant predictor aims to generate the favorable amino acid under the given fitness feature at a specific position. Moreover, pairwise dependencies between residues obtained by a Markov Random Field are also incorporated to promote reasonable variant generation. Experimental results on three viral protein datasets show that our EVPMM significantly outperforms baselines in most cases. Remarkably, our model also detects a potential mutational position 487 of SARS-CoV-2 which has not been reported by any other work, at which the generated variant may have an optimal binding with ACE2. In the future, we will conduct wet-lab experiment to rigorously verify the effectiveness of the virus variants generated by our model.

## Supplementary information

If your article has accompanying supplementary file/s please state so here.

Authors reporting data from electrophoretic gels and blots should supply the full unprocessed scans for key as part of their Supplementary information. This may be requested by the editorial team/s if it is missing.

Please refer to Journal-level guidance for any specific requirements.

## Acknowledgments

Acknowledgments are not compulsory. Where included they should be brief. Grant or contribution numbers may be acknowledged.

Please refer to Journal-level guidance for any specific requirements.

## Declarations

Some journals require declarations to be submitted in a standardised format. Please check the Instructions for Authors of the journal to which you are submitting to see if you need to complete this section. If yes, your manuscript must contain the following sections under the heading ‘Declarations’:

- Funding
- Conflict of interest/Competing interests (check journal-specific guidelines for which heading to use)
- Ethics approval
- Consent to participate
- Consent for publication
- Availability of data and materials
- Code availability
- Authors’ contributions

If any of the sections are not relevant to your manuscript, please include the heading and write ‘Not applicable’ for that section.

Editorial Policies for:

Springer journals and proceedings:

https://www.springer.com/gp/editorial-policies

Nature Portfolio journals:

https://www.nature.com/nature-research/editorial-policies

*Scientific Reports*:

https://www.nature.com/srep/journal-policies/editorial-policies

BMC journals:

https://www.biomedcentral.com/getpublished/editorial-policies

## Appendix A Section title of first appendix

An appendix contains supplementary information that is not an essential part of the text itself but which may be helpful in providing a more comprehensive understanding of the research problem or it is information that is too cumbersome to be included in the body of the paper.

